# Variability of 7K and 11K SomaScan plasma proteomics assays

**DOI:** 10.1101/2024.08.06.606813

**Authors:** Julián Candia, Giovanna Fantoni, Francheska Delgado-Peraza, Nader Shehadeh, Toshiko Tanaka, Ruin Moaddel, Keenan A. Walker, Luigi Ferrucci

**Affiliations:** Intramural Research Program, National Institute on Aging, National Institutes of Health, Baltimore, MD 21224, USA

## Abstract

SomaScan is an aptamer-based proteomics assay designed for the simultaneous measurement of thousands of human proteins with a broad range of endogenous concentrations. The 7K SomaScan assay has been recently expanded into the new 11K version. Following up on our previous assessment of the 7K assay, here we expand our work on technical replicates from donors enrolled in the Baltimore Longitudinal Study of Aging. By generating SomaScan data from a second batch of technical replicates in the 7K version, as well as additional intra- and inter-plate replicate measurements in the new 11K version using the same donor samples, this work provides useful precision benchmarks for the SomaScan user community. Beyond updating our previous technical assessment of the 7K assay with increased statistics, here we estimate inter-batch variability, we assess inter- and intra-plate variability in the new 11K assay, we compare the observed variability between the 7K and 11K assays (leveraging the use of overlapping pairs of technical replicates), and explore the potential effects of sample storage time (ranging from 2 to 30 years) in the assays’ precision.

## Introduction

SomaScan^1^ is a highly multiplexed, aptamer-based assay capable of simultaneously measuring thousands of human proteins broadly ranging from femto- to micro-molar concentrations. This technology relies on protein-capture SOMAmer (*Slow Offrate Modified Aptamer*) reagents,^2^ designed to optimize high affinity, slow off-rate, and high specificity to target proteins. These targets extensively cover major molecular functions including receptors, kinases, growth factors, and hormones, and span a diverse collection of secreted, intracellular, and extracellular proteins or domains. In recent years, SomaScan has increasingly been adopted as a powerful tool to discover biomarkers across a wide range of diseases and conditions, as well as to elucidate their biological underpinnings in proteomics and multi-omics studies.^3–10^

Concurrently with its wider adoption, SomaScan has expanded its proteome coverage by increasing the number of SOMAmers included in different versions of the assay, from roughly 800 SOMAmers in 2009, to 1,100 in 2012, 1,300 in 2015, 5,000 in 2018, 7,000 in 2020 and the most recent 11,000 protein assay released in November 2023.^11^ The first independent analysis of SomaScan normalization procedures and their technical variability was published by our teams at the U.S. National Institutes of Health (NIH) on the 1.1K and 1.3K assays, ^12^ later followed by reports from other laboratories^13–17^ and by an updated assessment based on the 7K assay.^18^ In the latter, we utilized inter-plate technical duplicates obtained from 102 human subjects enrolled in the Baltimore Longitudinal Study of Aging (BLSA), which allowed us to characterize different normalization procedures, evaluate assay variability, and provide detailed performance assessments.

In this work, we expand the scope of our previous study with the following aims: (i) to update the technical assessment of the 7K assay with the addition of a second batch of technical replicates, which allows us to increase statistics and to estimate inter-batch variability; (ii) to extend the assessment to the recently released 11K assay using inter- and intra-plate technical replicates; (iii) to compare the observed variability between the 7K and 11K assays, leveraging the use of overlapping pairs of technical replicates across the 7K and 11K assay versions; and (iv) to assess the potential relationship between technical variability and the time between sample collection and measurement over a time period spanning from 2 to 30 years.

## Experimental Section

### Study design

Fig. 1 shows details of the study design. Blood samples were collected from participants in the Baltimore Longitudinal Study of Aging^19^ (BLSA), a study of normative human aging established in 1958 and conducted by the National Institute on Aging, NIH. The study protocol was conducted in accordance with Declaration of Helsinki principles and was reviewed and approved by the Institutional Review Board of the NIH’s Intramural Research Program. Written informed consent was obtained from all participants. Following pre-processing using standard protocols, EDTA plasma vials were stored at *−*80*^◦^*C (panel (a)). After a period of 2-30 years of storage, samples were processed in SomaScan and analyzed (panel (b)). Samples were organized in 96*−*well (8 *×* 12) plates. In our design, we included intra- and inter-plate technical replicate pairs, as indicated in panel (c). In order to allow for a paired analysis across assay versions, all of the replicate samples run in the 11K platform were previously run in the 7K platform.

**Figure 1:**
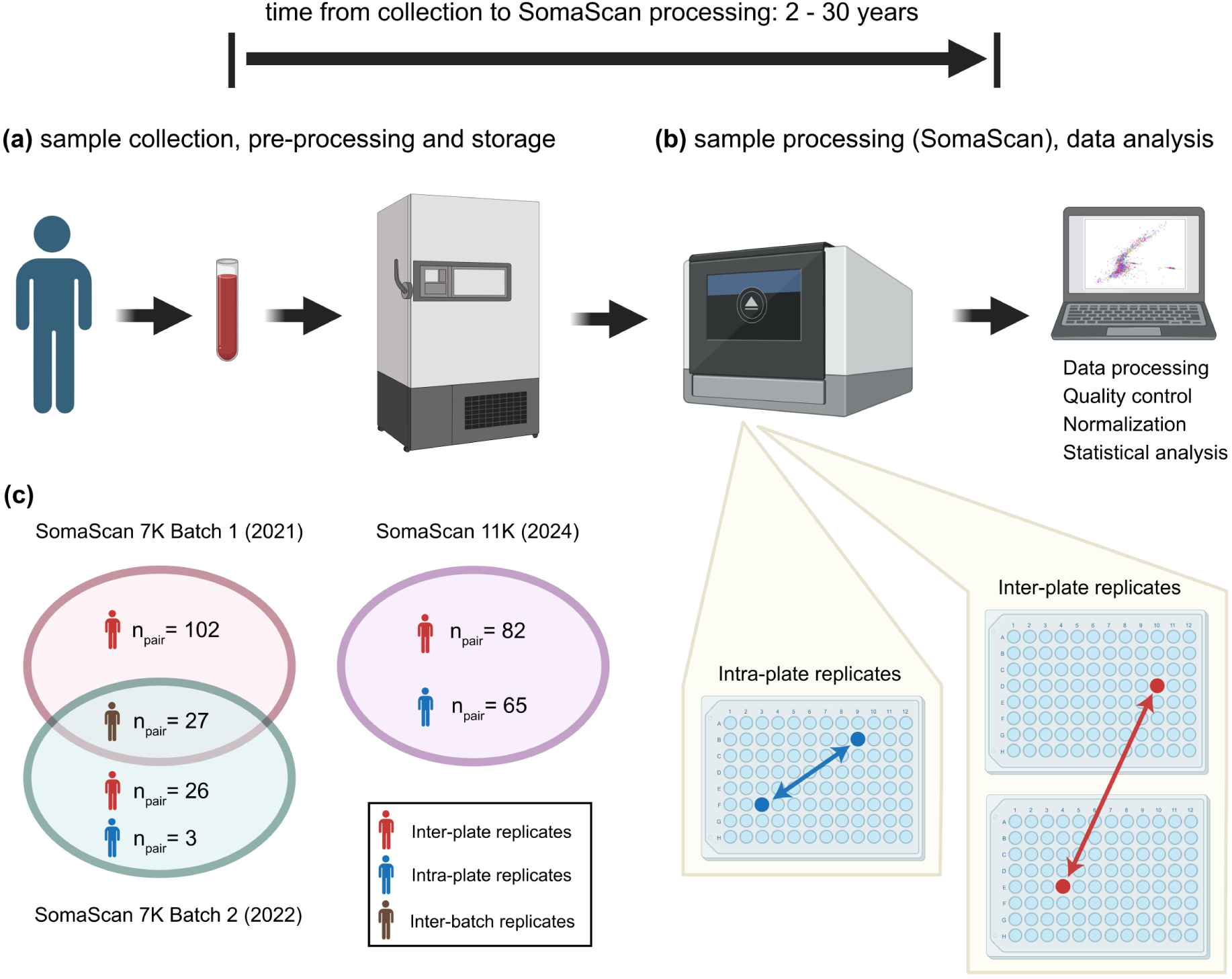
Schematic summary of the study. **(a)** Blood samples were collected from participants in the Baltimore Longitudinal Study of Aging (BLSA), then pre-processed and stored at *−*80*^◦^*C. **(b)** After a storage period of 2-30 years, samples were processed in SomaScan and analyzed. Some samples were run as intra- or inter-plate replicates. **(c)** Replicate pairs were run in two separate SomaScan 7K batches in years 2021 and 2022, respectively, followed by SomaScan 11K in year 2024. All of the replicate samples run in the 11K assay were previously run in the 7K assay.

### The SomaScan assay

Plasma proteomic profiles were characterized using the 7K (v4.1) and 11K (v5.0) SomaScan assays (SomaLogic, Inc.; Boulder, CO, USA). These assays consist of 7,289 and 10,776 SOMAmers targeting annotated human proteins, respectively. All of the SOMAmers in the 7K assay were included in the new 11K version. In order to cover a broad range of endogenous concentrations, SOMAmers are binned into different dilution groups, namely 20% (1:5) dilution for proteins typically observed in the femto- to pico-molar range (which comprise about 80% of all human protein SOMAmers in the assays), 0.5% (1:200) dilution for proteins typically present in nano-molar concentrations (slightly below 20% of human protein SOMAmers in the assays), and 0.005% (1:20,000) dilution for proteins in micro-molar concentrations (about 2 *−* 3% of human protein SOMAmers in the assays). The human plasma volume required is 55 *µ*L per sample.

The experimental procedure follows a sequence of steps, namely: (1) SOMAmers are synthesized with a fluorophore, photocleavable linker, and biotin; (2) diluted samples are incubated with dilution-specific SOMAmers bound to streptavidin beads; (3) unbound proteins are washed away, and bound proteins are tagged with biotin; (4) UV light breaks the photocleavable linker, releasing complexes back into solution; (5) non-specific complexes dissociate while specific complexes remain bound; (6) a polyanionic competitor is added to prevent rebinding of non-specific complexes; (7) biotinylated proteins (and bound SOMAmers) are captured on new streptavidin beads; and (8) after SOMAmers are released from the complexes by denaturing the proteins, fluorophores are measured following hybridization to complementary sequences on a microarray chip. The fluorescence intensity detected on the microarray, measured in RFU (*Relative Fluorescence Units*), is assumed to reflect the amount of available epitope in the original sample.

Raw data, as obtained after aggregation from slide-based hybridization microarrays, exhibit intra-plate nuisance variance due to differences in loading volume, leaks, washing conditions, etc, which is then compounded with inter-plate biases. In order to account for intra- and inter-plate effects, we apply a normalization procedure consisting of a sequence of steps, whose main elements are hybridization normalization, median signal normalization, plate-scale normalization, and inter-plate calibration (Table 1). Each of these steps generates scale factors at different levels: plate-specific, by SOMAmer dilution group, SOMAmer-specific, by sample type, and combinations thereof. Besides removing technical variability, these scale factors can be used as quality control flags at the plate-, sample-, and SOMAmer-levels. ^7^ For further details, see Refs. ^11,18^

**Table 1:** Summary of normalization steps.

| Step | Name | Abbreviation |
| --- | --- | --- |
| 0 | Raw | raw |
| 1 | Hybridization control normalization | hyb |
| 2 | Median signal normalization on calibrators | hyb.msnCal |
| 3 | Plate-scale normalization | hyb.msnCal.ps |
| 4 | Inter-plate calibration | hyb.msnCal.ps.cal |
| 5 | Median signal normalization on all sample types | hyb.msnCal.ps.cal.msnAll |

SOMAmers are uniquely identified by their “SeqId”, but the relation between SOMAmers and annotated proteins is not one-to-one. In the 11K assay, the 10,776 SOMAmers that target human proteins are mapped to 9,609 unique UniProt IDs, yielding a total of of 10,893 unique SOMAmer-UniProt ID pairs. Supplementary Data 1 provides the list of SOMAmer-UniProt ID pairs, along with protein annotations extracted from the UniProt Knowledge-base (UniProtKB), including length (number of amino acids in the canonical sequence), mass (molecular weight), pH dependence, Redox potential, temperature dependence, protein interactions, and subcellular location.

### Statistical methods

#### Coefficient of variation estimates from replicate pairs

A standard metric to characterize assay variability is the coefficient of variation (*CV*) defined as *CV* = (*σ/µ*) *×* 100%, i.e. the ratio between the standard deviation and the mean over a sufficiently large number of technical replicates. However, for technical duplicates, this definition is unstable because it relies on the calculation of standard deviations using just two data points per distribution (where each distribution describes replicate measurements from one subject-SOMAmer pair). Instead, we previously proposed a grid-search procedure that estimates *CV* from *n_dupl_* technical duplicates,^18^ as follows.

The theoretical relationship between the coefficient of variation and the expected fold change (*FC*) ratio between repeated measurements was derived by Reed, Lynn, and Meade. ^20^ Assuming that two replicate measurements, *RFU*_1_ and *RFU*_2_, are log-normally distributed random variables with the same mean and variance, we define the fold change *FC ≡ RFU*_1_*/RFU*_2_ to be restricted to *FC ≥* 1, without loss of generality. Then, the theoretical probability that these replicate measurements differ by a fold change equal or larger than *FC* is given by

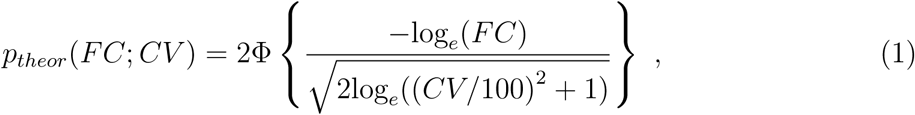

where Φ is the cumulative standard normal distribution function parametrized by *CV*.

For each SOMAmer, *FC* values obtained from replicate measurements define an empirical probability distribution, *p_emp_*(*FC*), whose similarity with *p_theor_*(*FC*; *CV*) may be evaluated via the non-parametric, one-sample Kolmogorov-Smirnov (KS) test. Therefore, we may explore a dense ensemble of theoretical distributions generated by a pseudo-continuous scanning of the *CV* parameter in Eq. (1) and evaluate the similarity between the target empirical distribution and each theoretical one as the *−log*_10_(p *−* value) derived from the one-sample KS test between *p_emp_*(*FC*) and *p_theor_*(*FC*; *CV*). Following this procedure, we thus estimate *CV* as the value that minimizes *−log*_10_(p *−* value) or, in other words, the *CV* value that generates the theoretical distribution most similar to the measured one.

### Data and Software Availability

Data analysis was performed in R v.4.4.0. Protein annotations from UniProtKB were extracted using the R package queryup^21^ v.1.0.5. Plots were generated using R packages RColorBrewer v.1.1-3, viridis v.0.6.5 and calibrate v.1.7.7. BioRender was used to prepare Fig. 1. Anonymized datasets and custom R source code used in our analyses are available on the Open Science Framework repository, https://osf.io/2b9qe (DOI 10.17605/OSF.IO/2B9QE).

## Results and Discussion

In order to explore the role of normalization in the 11K SomaScan assay, we used *n_dupl_* = 82 inter-plate technical duplicates to implement our grid-search procedure (described in the Methods section) to all human protein SOMAmers. The resulting *CV* density distributions are shown in Fig. 2(a) across all normalizations. Here, we observe that adding normalization steps shifts the distributions towards lower *CV* values. In greater detail, Fig. 2 (b-f) display cumulative distributions of the number of SOMAmers below a given *CV* for each normalization, which are consistent with previous reports on inter-plate variability in the 7K SomaScan assay.^18^ The estimated *CV* s for each SOMAmer and normalization step are provided for the updated 7K inter-plate estimates (Supplementary Data 2), as well as the new estimates for 7K intra-batch (Supplementary Data 3), 11K inter-plate (Supplementary Data 4) and 11K intra-plate (Supplementary Data 5). Unless stated otherwise, the remainder of this paper presents results for the fully normalized (“hyb.msnCal.ps.cal.msnAll”) data.

**Figure 2:**
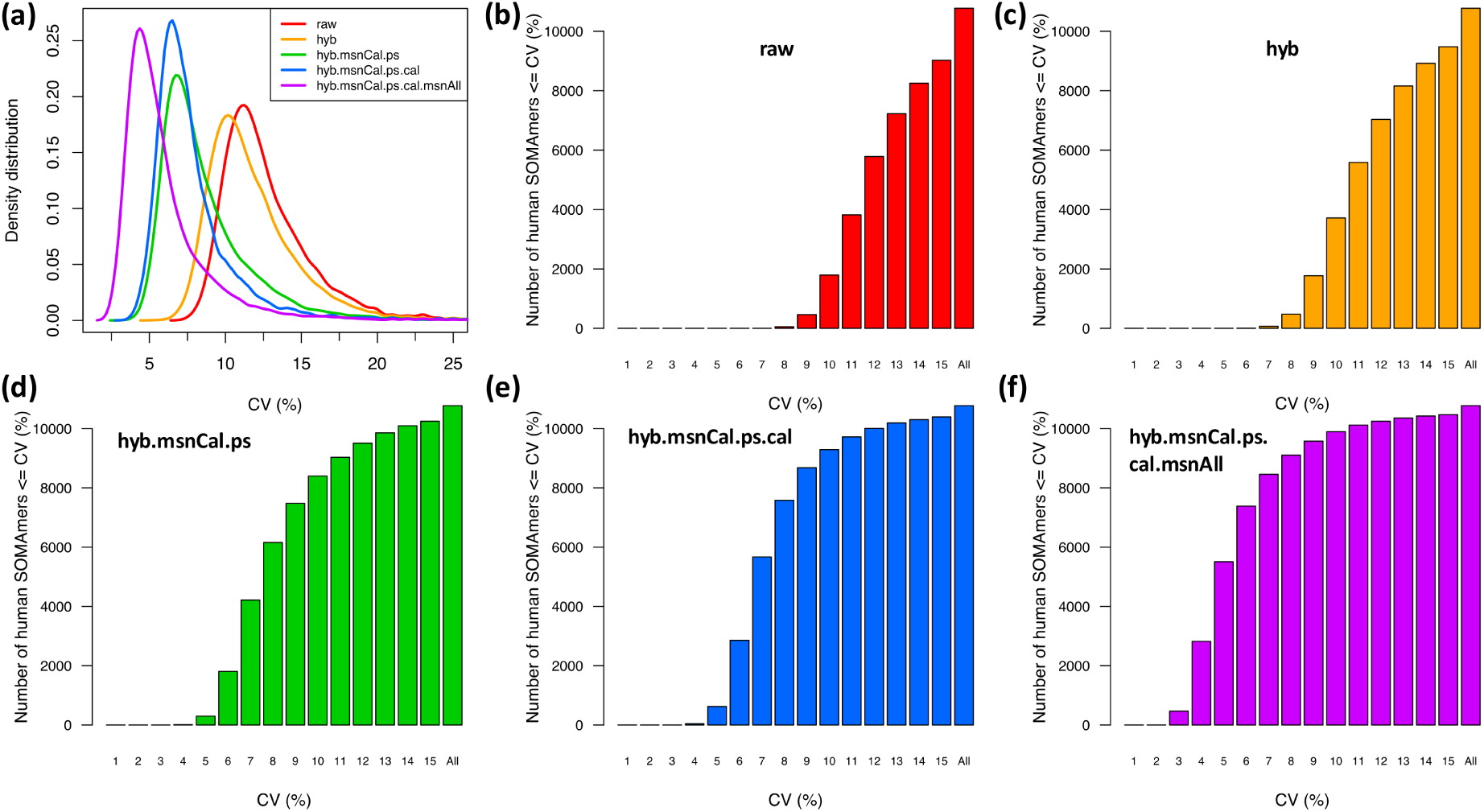
Distributions of coefficients of variation (CV). **(a)** Density distributions of *CV* across 10,776 human protein SOMAmers, obtained from inter-plate technical duplicates from 82 human subjects for different normalizations. **(b)-(f)** Cumulative distribution of the number of SOMAmers below a given *CV* for different normalizations, as indicated.

Table 2 shows a summary of the *CV* distributions by percentile for each case. On the one hand, we observe that inter-plate *CV* percentiles for 7K and 11K are similar (median *CV* = 4.5 and 5%, respectively). In greater detail, Fig. 3(a) shows that *CV* s comprising 7,289 human protein SOMAmers are strongly correlated. It is therefore reassuring that the overall precision of SomaScan has been maintained with the assay expansion from 7K to 11K. This is consistent with the findings of Rooney et al^22^ when comparing between 5K and 11K SomaScan versions. On the other hand, we observe that intra-plate variability is lower than inter-plate variability, as expected (median *CV* = 3 and 5%, respectively). As shown in Fig. 3(b), the correlation between intra- and inter-plate *CV* across 10,776 human protein SOMAmers remains strong, but the scatterplot cloud appears shifted in relation to the *y* = *x* reference line (dashes). Finally, we observe that the precision across batches is significantly degraded (despite the fact that we renormalized both batches together using the same set of calibrator replicates). Indeed, the median (intra-batch) inter-plate *CV* = 4.5% is found to increase to median inter-batch *CV* = 8. Correspondingly, Fig. 3(c) shows that *CV* s are correlated but shifted towards larger values in the inter-batch assessment relative to (intra-batch) inter-plate estimates.

**Table 2:** Summary of CV percentile distributions.

| CV Percentile | 7K Inter-plate | 7K Inter-batch | 11K Inter-plate | 11K Intra-plate |
| --- | --- | --- | --- | --- |
| 1% | 2.5 | 3 | 3 | 2 |
| 5% | 2.5 | 4 | 3.5 | 2 |
| 10% | 3 | 4.5 | 3.5 | 2.5 |
| 15% | 3 | 5 | 4 | 2.5 |
| 20% | 3.5 | 5.5 | 4 | 2.5 |
| 25% | 3.5 | 5.5 | 4 | 2.5 |
| **50%** | **4.5** | **8** | **5** | **3** |
| 75% | 6 | 11.5 | 7 | 4 |
| 80% | 6.5 | 13 | 7.5 | 4 |
| 85% | 7 | 15 | 8.5 | 4.5 |
| 90% | 8.5 | 18.5 | 9.5 | 5 |
| 95% | 11 | 23.6 | 12 | 5.5 |
| 99% | 23 | 38 | 27 | 9.5 |

**Figure 3:**
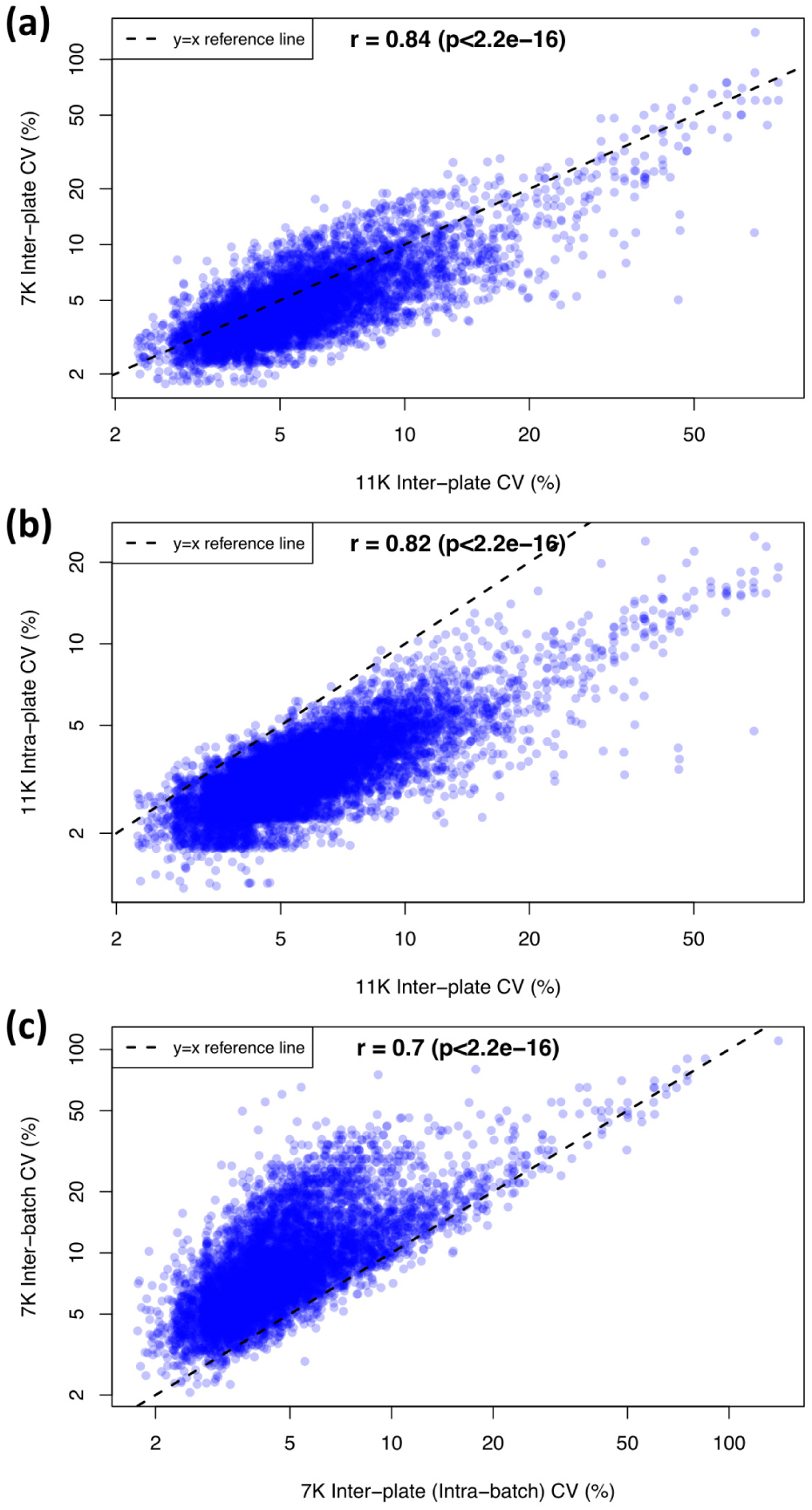
Comparison of coefficients of variation (CV). **(a)** 7K inter-plate versus 11K inter-plate. **(b)** 11K intra-plate versus 11K inter-plate. **(c)** 7K inter-batch versus 7K inter-plate (intra-batch). Pearson’s correlation estimates and p-values are shown. Data have been fully normalized.

One important design feature in our study is that all of the replicate samples run in the 11K assay were previously run in the 7K assay. More specifically, 74 inter-plate pairs were run in both assay versions, which allows us to perform a cross-version paired analysis of correlations across SOMAmers, as follows. For each replicate pair, we calculate the Spearman’s correlation estimate across all SOMAmers. Fig. 4(a) shows the correlations for all replicate pairs measured in the 7K (blue) and 11K (red) assay versions; matching samples across assays are joined by gray lines. The mean correlation estimate difference (defined as 11K minus 7K) is Δ*r* = *−*0.0051, indicating that the correlation is slightly higher in the 7K assay relative to the 11K assay, with a Wilcoxon paired test p *−* value = 0.0013. It should be noticed, however, that the 11K correlations in panel (a) were measured across all 10,776 human protein SOMAmers in the assay; in panel (b), we repeat the procedure but limiting 11K to the 7,289 human protein SOMAmers in common with the 7K assay, and observe again a small but significant negative difference between correlations in 11K and 7K. Since we notice five outlier samples, for which correlations in either assay appear unusually low, in panels (c-d) we repeated the analysis of panels (a-b) after removing these outliers. The conclusion, however, remains that the correlation across SOMAmers within inter-plate pairs appears to be slightly but significantly higher in the 7K SomaScan assay compared with the 11K assay version.

**Figure 4:**
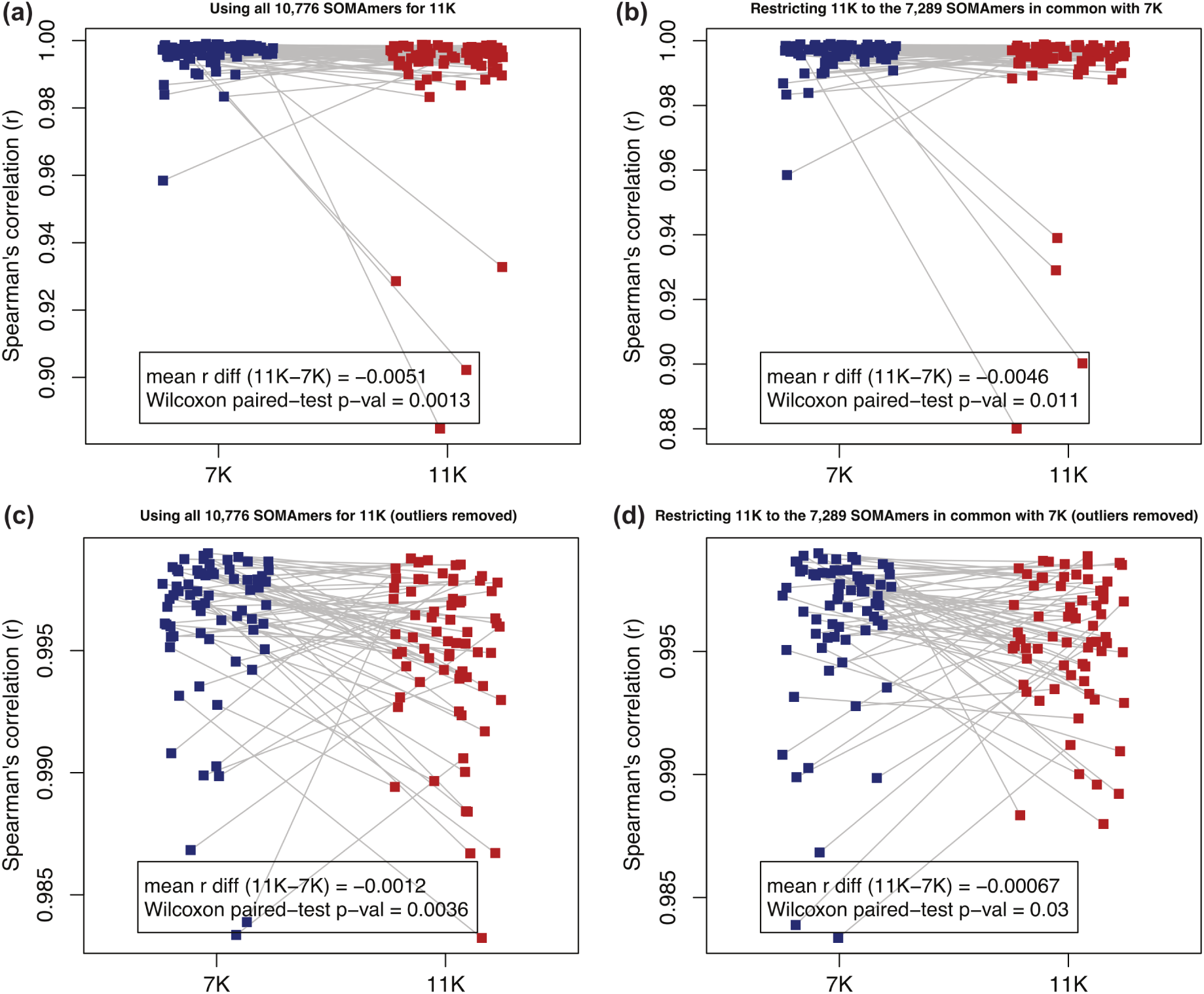
Inter-plate replicate correlations across SOMAmers: 11K vs 7K paired-sample comparison. For each inter-plate technical replicate, the Spearman’s correlation across SOMAmers is computed using the 7K (blue) and 11K (red) assays. Matching samples across assays are joined by gray lines. The mean difference between correlation estimates in the 11K assay relative to the 7K is shown, as well as the p-value from a Wilcoxon paired test. **(a)** Using 74 inter-plate technical replicates in each assay version (7,289 human protein SOMAmers in the 7K assay and 10,776 human protein SOMAmers in the 11K assay). **(b)** Using 74 inter-plate technical replicates in each assay version (but restricting the 11K assay to the 7,289 SOMAmers in common with 7K). **(c-d)** Same as panels **(a-b)**, but removing five outlier samples. Data have been fully normalized.

Fig. 5 shows the inter-plate *CV* as a function of the median signal-to-background ratio (*SBR*, defined as the median RFU of all samples divided by the median RFU of all buffer wells) for the 10,776 human protein SOMAmers in the 11K assay, colored by dilution group. The *SBR* estimates for each SOMAmer and normalization step are provided in Supplementary Data 6. The Spearman’s correlation between *CV* and *SBR* (*r* = 0.48) is primarily driven by the 20% dilution group (*r* = 0.51), which represents 80% of all human protein SOMAmers, followed by the 0.5% dilution group (*r* = 0.13), which represents 18% of all human protein SOMAmers. The third group, 0.005% dilution, comprises only 2% of all human protein SOMAmers and shows the opposite trend (*r* = *−*0.17). All correlation estimates are significant (*p <* 0.05). The *CV* median *±* MAD across SOMAmers are 5 *±* 1.5% (for all human protein SOMAmers as well as for the 20% dilution group), 6 *±* 2% (for the 0.5% dilution group), and 7 *±* 2% (for the 0.005% dilution group), respectively. The *SBR* median *±* MAD across SOMAmers are 7.8 *±* 7.1 (for all human protein SOMAmers), 6.4 *±* 5.1 (for the 20% dilution group), 21.9 *±* 22.3 (for the 0.5% dilution group), and 82.3 *±* 103.0 (for the 0.005% dilution group), respectively. Pairwise Wilcoxon test comparisons between dilution groups indicate that all these *CV* and *SBR* differences are statistically significant (adjusted p-value *<* 0.05). We have not observed any significant associations between *CV* or *SBR* and the protein characteristics described in Supplementary Data 1 (such as protein length, mass, or number of protein interactions).

**Figure 5:**
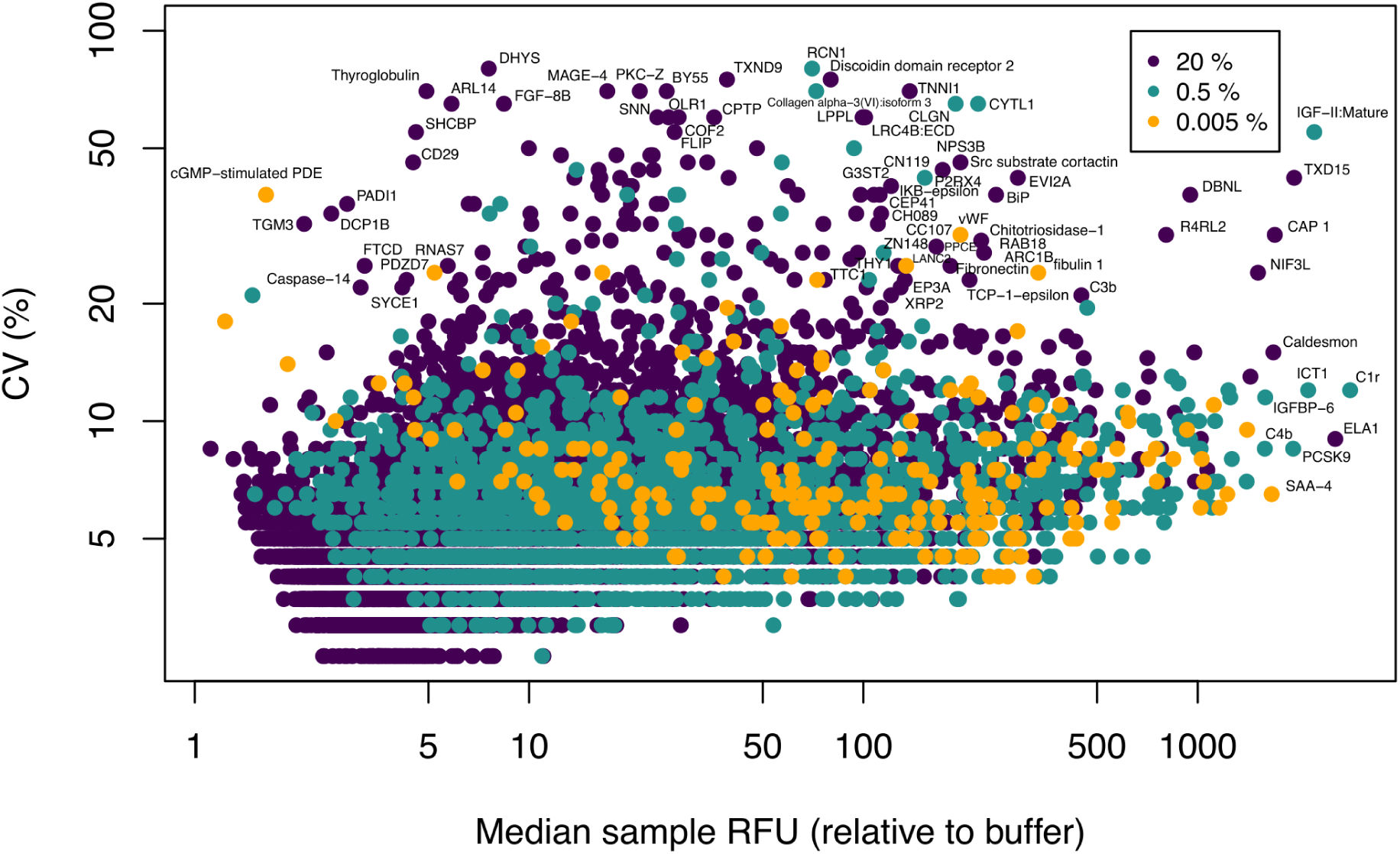
Brightness and variability of SOMAmers by dilution group. Inter-plate *CV* of 10,776 human protein SOMAmers as a function of the median signal-to-background ratio across experimental samples. SOMAmers are colored based on dilution: 20% (*n* = 8644), 0.5% (*n* = 1911), and 0.005% (*n* = 221). Data have been fully normalized.

Finally, we explored the association between technical variability and sample storage time. For the 7K SomaScan assay, 128 inter-plate pairs were measured between 2 and 28 years after the time of sample collection, pre-processing and storage. For each SOMAmer, we calculated the correlation between the inter-plate replicate fold change and the sample storage time (Supplementary Data 7). This is summarized in Fig. 6(a), which shows the p-value of the Spearman’s correlation (transformed as *−log*_10_(p *−* value) on the y-axis) as a function of the SOMAmer *CV* (on the x-axis). The dashed line shows the FDR-adjusted significance threshold *q* = 0.05. We observe that only seven proteins appear above the significance threshold. All of them are positively correlated, as shown by Fig. 6(b), which is consistent with the interpretation that protein degradation is responsible for increased variability over time. These effects are shown in greater detail in panels (c-d) for the top two proteins, namely collagen type VI alpha 1 chain (COL6A1) and tyrosinase related protein 1 (TYRP1/gp75), respectively. Other proteins with significant degradation are related to plasma and coagulation. In contrast, two proteins reported to show the strongest association with human aging,^5^ namely pleiotrophin (PTN) and macrophage inhibitory cytokine-1 (MIC-1/GDF15), show mild to insignificant degradation effects with storage time, as shown in panels (e-f), respectively. These observations have important consequences for the interpretation of findings from longitudinal studies such as BLSA, in which samples are typically stored for many years (and even decades) before they are thawed and processed to generate data.

**Figure 6:**
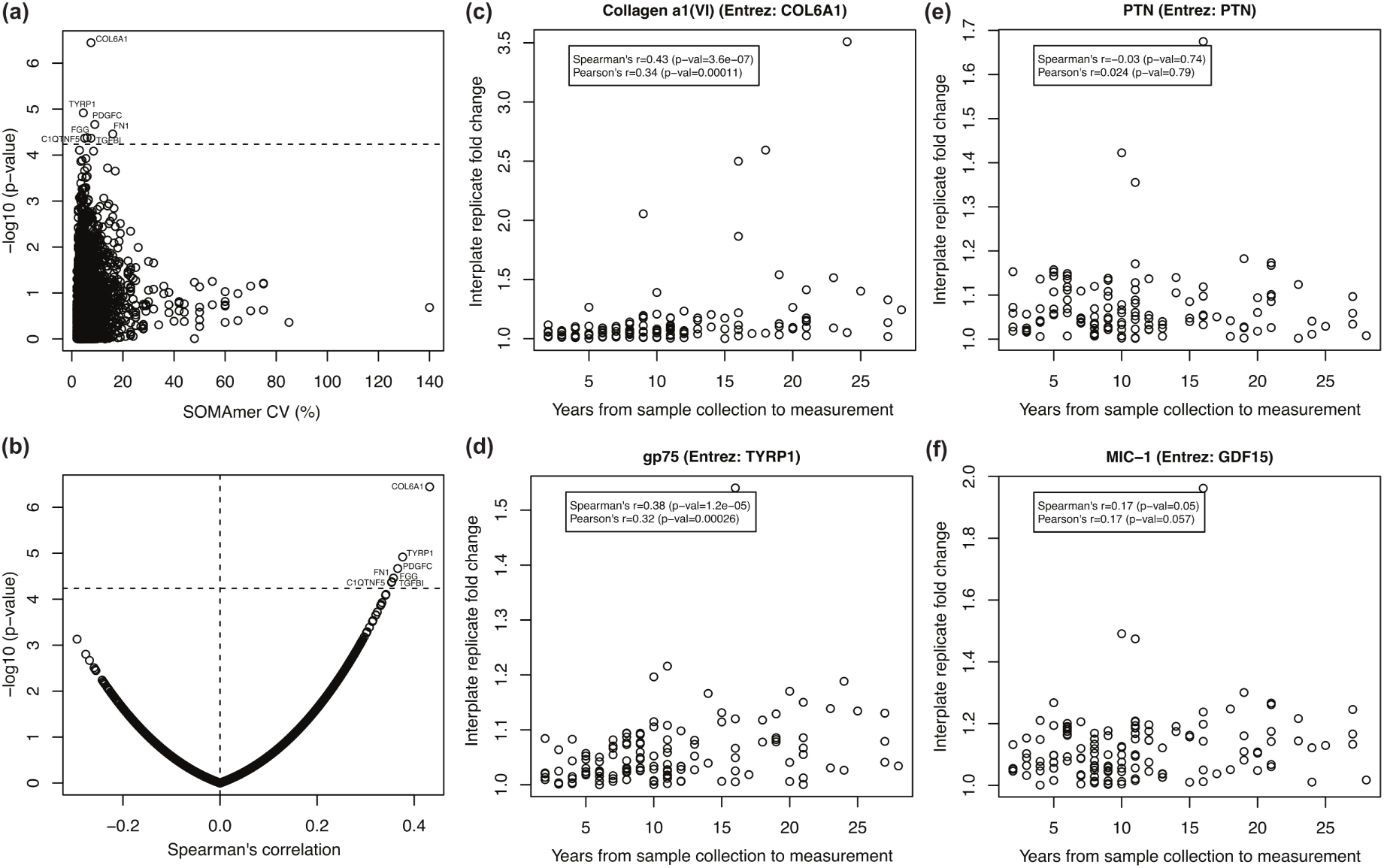
Association between technical variability and storage time in the 7K SomaScan assay. **(a)** Significance of the association between SOMAmer variability and storage time (y-axis) as a function of SOMAmer *CV* (x-axis). The dashed line shows the FDR-adjusted significance threshold *q* = 0.05. **(b)** Significance of the association between SOMAmer variability and storage time (y-axis) as a function of the Spearman’s correlation (x-axis). **(c-f)** Inter-plate replicate fold change (*FC*) as a function of storage time. Spearman’s and Pearson’s correlation estimates and p-values are also shown. Panels **(c-d)** show the two SOMAmers with the strongest association (COL6A1 and TYRP1), whereas panels **(e-f)** showcase the two SOMAmers previously reported to exhibit the strongest association with aging (PTN and GDF15), which do not show significant effects with storage time. Data have been fully normalized.

Similarly, Fig. 7 shows the association between technical variability and sample storage time for the 11K SomaScan assay, based on 82 inter-plate pairs stored for 5 to 30 years (Supplementary Data 8). None of the SOMAmers shows a significant degradation (using the *q* = 0.05 threshold after FDR adjustment), as shown in panel (a), although the top SOMAmers appear biased towards positive correlations (panel (b)), indicating that protein degradation may be occurring on these SOMAmer targets to a moderate degree. Data for the top two proteins, serine protease 27 (marapsin/PRSS27) and betacellulin (BTC), is shown in panels (c-d), respectively. Aging-associated proteins PTN and GDF15, shown in panels (e-f), display no significant degradation effects due to storage time. Many of the samples used in this study have not been aliquoted for single-use thawing, but we lack full historical records to track which of them may have been frozen and thawed multiple times. A separate study on the impact caused by repeated freeze-thaw cycles in 11K SomaScan quantification shows only 4% of affected SOMAmers; ^23^ the overlap between affected probes by storage (690 SOMAmers obtained from Spearman’s p-value < 0.05, Supplementary Data 8) and affected probes by freeze-thaw cycles (466 SOMAmers with p-value < 0.05 derived from mixed-effects models^23^) is not statistically significant (31 overlapping SOMAmers, Fisher’s exact test p *−* value = 0.77).

**Figure 7:**
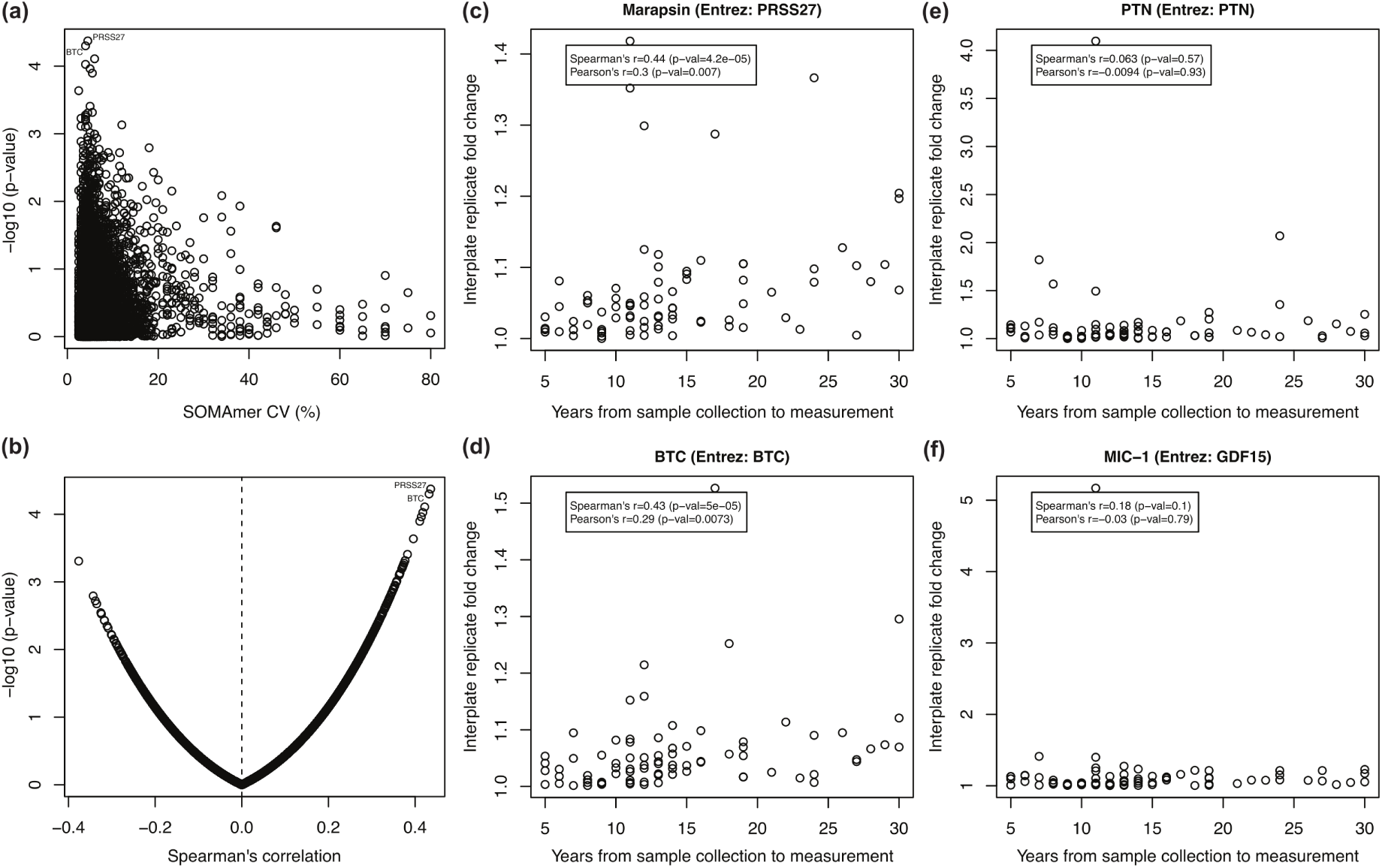
Association between technical variability and storage time in the 11K SomaScan assay. **(a)** Significance of the association between SOMAmer variability and storage time (y-axis) as a function of SOMAmer *CV* (x-axis). None of the observations reaches the FDR-adjusted significance threshold *q* = 0.05. **(b)** Significance of the association between SOMAmer variability and storage time (y-axis) as a function of the Spearman’s correlation (x-axis). **(c-f)** Inter-plate replicate fold change (*FC*) as a function of storage time. Spearman’s and Pearson’s correlation estimates and p-values are also shown. Panels **(c-d)** show the two SOMAmers with the strongest association (PRSS27 and BTC), whereas panels **(e-f)** showcase the two SOMAmers previously reported to exhibit the strongest association with aging (PTN and GDF15), which do not show significant effects with storage time. Data have been fully normalized.

## Conclusions

Similarly to previous versions of the SomaScan assay,^12,18^ the standard normalization procedure of 11K SomaScan data appears to operate as expected. While the median inter-plate *CV* of raw RFU counts was *CV_med_* = 12%, successive normalization steps reduced it after hybridization (*CV_med_* = 11%), plate-scale normalization (*CV_med_* = 8%), inter-plate calibration (*CV_med_* = 7%), and the fully-normalized data including median signal normalization applied on all sample types (*CV_med_* = 5%). It is reassuring to observe no precision degradation in the assay expansion from the 7K version to the 11K version. Moreover, we observed a strong correlation of *CV* estimates across human protein SOMAmers between 7K and 11K.

Our findings have implications for the plate design of SomaScan studies. Intra-plate variability (*CV_med_* = 3%) is significantly lower than inter-plate variability (*CV_med_* = 5%). Therefore, in order to prevent the occurrence of inter-plate effects in longitudinal studies, it is recommended to keep all longitudinal samples from the same subject together in the same plate and randomize across plates based on relevant covariates (e.g. age and sex), which can be implemented with tools such as the R package OlinkAnalyze. Inter-batch variability (*CV_med_* = 8%) is significantly higher than (intra-batch) inter-plate variability (*CV_med_* = 5%). The use of calibrators to account for inter-plate effects does not suffice to remove additional cross-batch effects. It is thus recommended that all samples in a study be run, whenever possible, within the same batch. Additional procedures for cross-batch correction may include the use of bridge samples to be added to runs across batches, as well as the exploration of additional normalization procedures. Regarding the latter, bioinformatic tools developed to assess and remove batch effects for other omics may be useful in the context of SomaScan data processing, for instance: ComBat, ^24^ implemented in the sva R package,^25^ a very well established method for batch correction in microarray and RNA-seq data; guided PCA;^26^ and multi-MA normalization, ^27^ among others. This important topic certainly deserves further investigation.

Using fully normalized data, we find that correlations between inter-plate replicate pairs across all human protein SOMAmers are close to 1 for both 7K and 11K SomaScan assay versions. However, a paired comparison (allowed by the fact that we used overlapping pairs of technical replicates across both assay versions) reveals that the correlation is slightly but significantly higher in the 7K SomaScan assay compared with the 11K assay version. Both assay versions exhibit a remarkable sensitivity, with 98% of the measured human proteins appearing *>* 2*−*fold brighter in human plasma samples compared to buffer wells.

Finally, we explored the important question of the possible association between technical variability and sample storage time. If increased protein degradation was found to significantly affect the assay’s precision, that would impose limits on the use of older samples stored in specimen banks, with particularly strong implications on the proteomic interrogation of long-established longitudinal studies such as the BLSA. We observed moderate degradation that appears to be protein-specific and not significantly associated with SOMAmers found to be affected by repeated freeze-thaw cycles;^23^ hence, devising a correction process to account for this effect may be complex. Fortunately, we found that the variability increase in samples stored in a time frame from 2 to 30 years was not significant for most SOMAmers.

As SomaScan becomes more widely adopted and utilized as a state-of-the-art tool for proteomics discovery, we hope that this work will serve as a valuable technical reference and resource for the growing SomaScan user community.

## Acknowledgement

This research was supported entirely by the Intramural Research Program of the National Institute on Aging, NIH.

## Supporting Information Available

The following files are available online:

- Supplementary Data 1: Protein annotations extracted from the UniProt Knowledge-base (UniProtKB).
- Supplementary Data 2: 7K SomaScan inter-plate coefficients of variation.
- Supplementary Data 3: 7K SomaScan inter-batch coefficients of variation.
- Supplementary Data 4: 11K SomaScan inter-plate coefficients of variation.
- Supplementary Data 5: 11K SomaScan intra-plate coefficients of variation.
- Supplementary Data 6: 11K SomaScan signal-to-background ratio estimates.
- Supplementary Data 7: 7K SomaScan correlation between inter-plate replicate fold change and sample storage time.
- Supplementary Data 8: 11K SomaScan correlation between inter-plate replicate fold change and sample storage time.

